# A leg model based on anatomical landmarks to study 3D joint kinematics of walking in *Drosophila melanogaster*

**DOI:** 10.1101/2024.04.04.588055

**Authors:** Moritz Haustein, Alexander Blanke, Till Bockemühl, Ansgar Büschges

**Author notes:** **Correspondence:** Ansgar Büschges. Shared senior authorship (Till Bockemühl, Ansgar Büschges).

## Abstract

Walking is the most common form of how animals move on land. The model organism *Drosophila melanogaster* has become increasingly popular for studying how the nervous system controls behavior in general and walking in particular. Despite recent advances in tracking and modeling leg movements of walking *Drosophila* in 3D, there are still gaps in knowledge about the biomechanics of leg joints due to the tiny size of fruit flies. For instance, the natural alignment of joint rotational axes was largely neglected in previous kinematic analyses. In this study we therefore present a detailed kinematic leg model in which not only the segment lengths but also the main rotational axes of the joints were derived from anatomical landmarks, namely the joint condyles. Our model with natural oblique joint axes is able to adapt to the 3D leg postures of straight and forward walking fruit flies with high accuracy. When we compared our model to an orthogonalized version, we observed that our model showed a smaller error as well as differences in the used range of motion (ROM), highlighting the advantages of modeling natural rotational axes alignment for the study of joint kinematics. We further found that the kinematic profiles of front, middle, and hind legs differed in the number of required degrees of freedom as well as their contributions to stepping, time courses of joint angles, and ROM. Our findings provide deeper insights into the joint kinematics of walking in *Drosophila*, and, additionally, help to develop dynamical, musculoskeletal, and neuromechanical simulations.

## 1 Introduction

Animals exhibit a rich repertoire of locomotive behaviors. In the context of legged locomotion, i.e. walking, animals can change their speed and heading direction, traverse diverse substrates, or can even compensate for the loss of a leg. This versatility emerges from the fact that biological limbs typically have more joints and/or more degrees of freedom (DOFs), i.e. independent directions of motions, than strictly required for any single movement task (Full and Koditschek, 1999; Bernstein and Latash, 2021). Consequently, detailed kinematic analyses are required to understand the demands on the underlying motor control system.

The neurobiology of walking is frequently studied in insects, because on the one hand insects and vertebrates follow many of the same general principles of locomotion (Pearson, 1993; Duysens et al., 2000; Büschges, 2005), while on the other hand their relatively simple nervous systems greatly facilitate the investigation of the underlying neuronal activity (Bidaye et al., 2018). Particularly, the fruit fly *Drosophila melanogaster* is an outstanding model organism for deciphering the motor control of walking, as it offers enormous potential for linking anatomy, physiology, and behavior: an ever-expanding genetic toolbox is available (Venken et al., 2011) for tracking individual neurons and recording or manipulating their activity in constrained preparations as well as in behaving flies (e.g. Chen et al., 2018; Mamiya et al., 2018; Azevedo et al., 2020; Bidaye et al., 2020; Feng et al., 2020; Chockley et al., 2022; Hermans et al., 2022). Furthermore, ongoing work to map the entire connectome of the brain (Scheffer et al., 2020) and the ventral nerve cord (Phelps et al., 2021; Takemura et al., 2023) is aiding in unravelling the neuronal circuits involved in various aspects of motor control on the synaptic and circuit level.

However, *Drosophila*’s tiny size and capability for relatively fast movements hampered 3D motion capture at the level of leg joints until recent advances in deep learning pose estimation algorithms (Günel et al., 2019; Karashchuk et al., 2021). Although 3D motion capture of the leg joints from walking fruit flies is already informative and useful in many experimental settings, pose estimation algorithms can typically provide only one tracked point per joint. Thus, joint angles from ball-and-socket joints with three DOFs, such as the thorax-coxa joint in the insect leg, must be estimated by using either projections of the coxa onto individual body planes (Lobato-Rios et al., 2022) or Euler angle sequences obtained by fitting a body coordinate system to the thorax-coxa joint (Bender et al., 2010; Karashchuk et al., 2021). These methods might, however, reflect only equivalent paths of motion rather than the true biological movements of limbs around actual joint axes (Woltring, 1991; Crawford et al., 1999). Another limitation of current pose estimation algorithms is that they are less accurate in capturing rotations about the longitudinal axis of limb segments (Ceseracciu et al., 2014) and locations of joint centers (Needham et al., 2021) compared to traditional marker-based motion capture approaches.

These issues can be overcome by using a multibody kinematics optimization strategy, which is widely used to compensate for soft tissue artefacts or building musculoskeletal models in human motion studies (Lu and O’Connor, 1999; Begon et al., 2018). For this, a 3D kinematic body model with rigid segments and well-defined joint DOFs is fitted to motion captured joint positions by applying a global optimization algorithm. Since in this approach the lengths of the segments and the joint rotational axes are preserved during movements, the postures resulting from these models can more accurately represent the actual alignment of body parts, such as legs, than 3D reconstructions based solely on singular joint positions. In fact, several 3D kinematic leg models have successfully been implemented in the last decades to study walking in stick insects (Zakotnik et al., 2004; Theunissen and Dürr, 2013; Dallmann et al., 2016), crickets (Petrou and Webb, 2012), ants (Arroyave-Tobon et al., 2022), and recently also in *Drosophila* (Goldsmith et al., 2022; Lobato-Rios et al., 2022). A challenge in developing accurate kinematic models is to define the correct parameters, for instance for segment lengths, number of DOFs of each joint, and the orientation of their rotational axes, as these design decisions directly affect the joint angles that will be obtained (Begon et al., 2018). Due to the small size of insects, most insect leg models had to rely on assumptions from kinematic studies and/or anatomical descriptions from morphological studies. Nowadays, improved models can be created based on extremely detailed 3D body reconstructions of insects obtained from nano- and micro-computed tomography (µCT) data (Blanke et al., 2017; Arroyave-Tobon et al., 2022; Dinges et al., 2022; Lobato-Rios et al., 2022). However, in most current kinematic leg models of insects, including *Drosophila*, the joint rotational axes were positioned generically perpendicular to the leg segments, an assumption that need not be true. Rotational axes of biological joints commonly show oblique orientations (Krause and Dürr, 2004; Rubenson et al., 2007; Frund et al., 2022), which should consequently affect not only the joint kinematics, but also other biomechanical aspects such as joint torques, the required muscle activation pattern, and, ultimately, the underlying neuronal control.

In this work, we therefore aimed to create a kinematic leg model for *Drosophila* in which the joint axes were aligned using the positions of the joint condyles as anatomical landmarks. This resulted in an oblique main axis of rotation of the individual joints. Afterwards, we used an inverse kinematic solver to fit our model to motion captured leg postures of straight, forward walking fruit flies and analyzed the resulting joint kinematics. To explore the importance of axis alignment, we compared our model to an alternative version in which the rotational axes were aligned orthogonally to the leg segments and found that our model with oblique joint axes showed a smaller error and used different ranges of motion (ROMs) for many joint DOFs. Moreover, we found that the front, middle, and hind legs have distinct kinematic profiles in terms of joint angles, ROMs, and contributions of each joint DOF. Our findings therefore not only provide relevant biomechanical insights into walking in *Drosophila*, but can also guide the development and improvement of more sophisticated models for dynamical, musculoskeletal and neuromechanical simulations.

## 2 Material and methods

### 2.1 Experimental animals

To robustly induce forward walking, we used Bolt-GAL4>UAS-CsChrimson *Drosophila melanogaster* flies (Bidaye et al., 2020; Bolt-GAL4 kindly provided by Dr. Salil Bidaye, UAS-CsChrimson BDSC-#55134) for all experiments. Animals were reared on a standard yeast-based medium (Backhaus et al., 1984) at 25°C and 65% humidity in a 12h:12h day:night cycle. All data were obtained from experiments with 3-8 days post-eclosion males (N=7) and females (N=5). Prior to experiments, the animals were kept in the dark for at least three days in fresh vials in which the food was soaked with 50 µL of a 100 mmol L^−1^ all-trans-Retinal solution (R2500; Sigma-Aldrich, RRID:SCR_008988).

### 2.2 Motion-capture setup

To capture joint movements, tethered flies walked stationarily on a spherical treadmill setup as described in detail previously (Berendes et al., 2016; Szczecinski et al., 2018). In brief, the animals were cold-anesthetized and a L-shaped copper wire (Ø: 0.15 mm) was attached to the dorsal side of the thorax with a small drop of light-curing adhesive (ESPE Sinfony, 3M ESPE AG, Seefeld, Germany), which was cured with a blue laser (460 nm). The tethered flies were positioned on top of a polypropylene ball (Ø: 6 mm; Spherotech GmbH, Fulda, Germany) using a 3D-micromanipulator. As the ball was air-suspended, it could be moved freely by the flies around its three axes of rotation. To promote walking behavior, the flies were centered on the ball so that their lateral and vertical orientations were straight relative to the ball surface and their ground clearance was adjusted accordingly. A red laser (658 nm) targeting the animal’s head was used to optogenetically activate the Bolt protocerebral neurons (BPNs) in the brain of the Bolt-GAL4>UAS-CsChrimson flies by opening the light-gated cation channel CsChrimson (Klapoetke et al., 2014). As BPNs are associated with the initiation and maintenance of fast forward walking (Bidaye et al., 2020), this allowed us to increase the number of walking sequences (**supplemental video 1**).

Movements of the ball around its three axes of rotation were measured at 50 Hz by two optical sensors (ADNS-9500; Avago Technologies, San Jose, USA) pointing at the ball’s equator and placed orthogonally to each other (**Figure 1A**). This allowed for the calculation of the global rotation of the ball and subsequent reconstruction of the virtual walking trajectory and forward speed of the animals (Seelig et al., 2010; Berendes et al., 2016; Szczecinski et al., 2018). We recorded only straight walking sequences at a relative constant walking speed (mean: 14.7 ± 4.0 mm per second, n=2250 steps).

**Figure 1.**
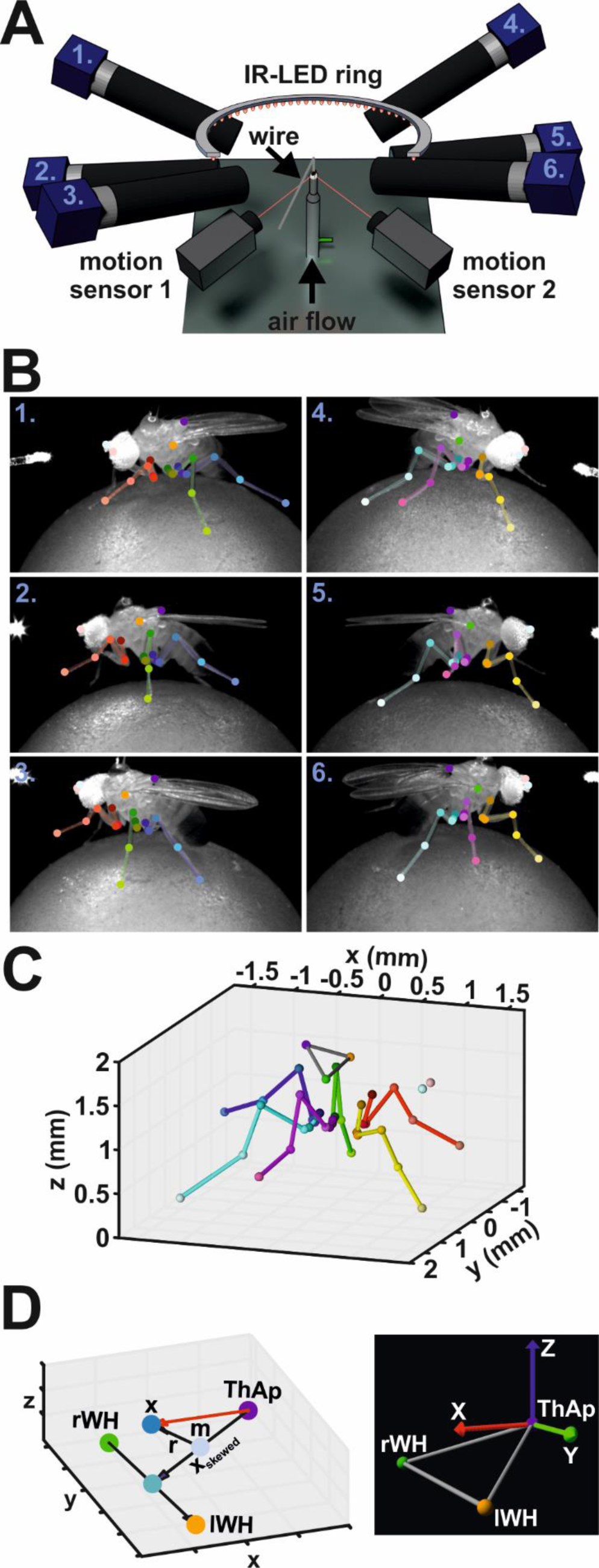
Motion Capture Setup. (**A**) Schematic illustration of the motion capture system. (**B**) Annotated video frames for all six camera views at the same time point. Numbers indicate the camera in A. (**C**) Corresponding 3D reconstruction derived from annotations in B. (**D**) Calculation of body coordinate system. Left panel: graph displays the vectors used to calculate the body coordinate system. Right panel: example image of a body coordinate system and the body reference keypoints used in the model visualizations.

Leg movements were recorded with six synchronized high-speed cameras (acA1300-200um, Basler AG, Ahrensburg, Germany) equipped with 50 mm lenses (LM50JC1MS, Kowa Optical Products Co. Ltd., Nagoya, Japan). For subsequent 3D reconstruction, cameras were placed around the animal such that multi-view images were obtained from either body side with a front, side, and hind aspect (**Figure 1A, supplemental video 1**). A supplementary camera (acA1300-200um) recording the scene from above was used for proper positioning of the animal on the ball and for camera calibration, but was not used for data acquisition. Illumination was achieved by a custom-built infrared light-emitting diode (IR-LED, wavelength 880 nm) ring. The synchronization of cameras, motion sensors, and IR-LED ring were accomplished by a custom-built controller device. Videos were acquired at 400 Hz, a resolution of 896 x 540 pixels (width x height, average field of vision: 5.2 mm x 3.2 mm, average spatial resolution of 5.9 ± 0.4 µm per pixel), and an exposure time of 500 µs. A custom-written non-linear contrast enhancement function was applied to the videos to improve visibility of the leg joints and the tarsus tip for subsequent tracking. Videos were compressed using the FFmpeg library (version N-93252-gf948082e5f). The compression settings used here (codec: libx264, constant rate factor: 12, preset: ultrafast) resulted in a file size reduction of about 90% while maintaining over 98% of the video quality as evaluated by structural similarity index measure (SSIM, Wang et al. 2004). Camera control was implemented based on the Harvester image acquisition library (version 1.3.1, available at: https://github.com/genicam/harvesters). All custom-made devices were built by the Electronics workshop of the Institute of Zoology, University of Cologne.

### 2.3 Automated tracking of leg and body keypoints

Detection of keypoints in the videos was performed with the DeepLabCut toolbox (version 2.2rc3; Mathis et al. 2018; Nath et al. 2019). For each leg, we tracked six keypoints: the thorax-coxa joint (ThCx), the coxa-trochanter joint (CxTr), the trochanter-femur joint (TrFe), the femur-tibia joint (FeTi), the tibia-tarsus joint (TiTar), and the tip of the tarsus (Tar) (**Figure 1B**). In addition, the posterior scutellum apex on the thorax (ThAp), the wing hinges (left, lWH; right, rWH), and the antennae (left, lAnt; right, rAnt) were tracked as body reference keypoints. We trained three independent ResNet-50 networks for videos from cameras having the same viewpoint for both sides of the body, i.e. a single network was trained for either both front, both side, and both hind camera views. The training sets for each network were generated by manual annotation of walking sequences of six flies (three males, three females) and contained a total number of 628, 755, 753 video frames for the front, side, hind networks, respectively. When keypoint occlusion occurred in a frame, we added position estimates to the training data to obtain complete positional sets for subsequent 3D reconstruction. Although this was a potential source of inaccuracies of tracked positions, the resulting impact was considered minor because occlusions occurred only transiently by movements of the leg segments in a relatively small area. Furthermore, they were mainly observed in the proximal joints, i.e. ThCx, CxTr, TrFe, whose positions did not change much during walking compared to the more distal joints. The training data sets were expanded by data augmentation techniques, i.e. cropping, rotation, brightness, blurring, and scaling, using the default in-built augmentation algorithm of DeepLabCut. All resulting keypoint predictions for experimental videos were inspected visually and erroneous keypoint predictions were corrected manually (5.6 % of all tracked keypoints). These manual annotations of keypoints and corrections were carried out with a custom-written graphical user interface.

### 2.4 3D reconstruction of tracked keypoints

Because each body keypoint was tracked from three different synchronized camera perspectives, their corresponding 3D position can be reconstructed by triangulation (**Figure 1C, supplemental video 1**). For this, we modeled each camera as pinhole camera with lens distortion (Hartley and Zisserman, 2004; Günel et al., 2019; Karashchuk et al., 2021). To determine the camera parameters, we calibrated the camera setup with a custom-made checkerboard pattern with 7 x 6 squares (square dimension: 399 x 399 µm). The checkerboard pattern was developed on a photographic slide (Gerstenberg Atelier für Visuelle Medien, Templin, Germany), cut out, and clamped in a custom-made metal frame to flatten the pattern. Images from the checkerboard pattern were acquired at full camera resolution (1280 x 1024 pixels). Camera calibration were performed with the OpenCV software library (Bradski, 2000). The corners of the pattern were detected on each image at sub-pixel level. Afterwards, the intrinsic parameters and lens distortion parameters (three and two coefficients for radial and tangential distortion, respectively; G. R. Bradski and Kaehler 2008) were determined for each camera (n > 60 images per camera, average reprojection error: 0.37 ± 0.07 pixels). The principal point was fixed to the center of the calibration images (x=640, y=512). Determination of camera extrinsic parameters based on an iterative stereo calibration procedure. For this, the checkerboard was positioned in a way that it was imaged simultaneously by two adjacent cameras (n > 20 per camera pair) and the differences between the corner positions due to the different viewing angles were used to derived the rotation matrix and translation vector required to match the reference camera to the other camera (average reprojection error: 0.76 ± 0.19 pixels). The order of camera pairs for each body side was front-to-side cameras and side-to-hind cameras. The position and orientation of the camera recording the scene from above (see **2.2**) served to define the global coordinate system and the results of the pairwise stereo calibration processes were used to determine the position and orientation for each camera in the global coordinate system.

Reconstruction of 3D positions of leg and body keypoints was achieved by triangulation (Hartley and Zisserman, 2004; Günel et al., 2019; Karashchuk et al., 2021). For this, a linear system of equations based on the corresponding image coordinates of the keypoints and the projection matrices of the cameras was solved by using singular value decomposition (SVD). To compensate for lens distortion, image coordinates were corrected prior to the SVD procedure by using the inversed distortion function of the respective camera based on the coefficients obtained during calibration. Code for SVD procedure was based on ‘Python Projective Camera Model’ module (author: Matej Smid, https://github.com/smidm/camera.py)

A body coordinate system was created for each fly to describe the 3D positions of keypoints in relation to the body of the animals and to adjust the kinematic leg model to the fly’s legs (**Figure 1D**, **Eqs. 1**). The origin of the body coordinate system was defined by the ThAp position. The y-axis was derived from the positions of the wing hinges (lWH, rWH) and pointed towards the left body side. To obtain an x-axis pointing towards the anterior direction of the body, the vector between the ThAp and the midpoint between the wing hinges was calculated at first, but the resulting vector was skewed (**x_skewed_**) because the wing hinges were situated ventrally in relation to the ThAp. However, given that **x_skewed_** is the hypotenuse of a right triangle, the adjacent leg lies on the desired x-axis. Therefore, we determined the intersection point of the adjacent and the opposite legs of the triangle by calculating the reflecting vector **r** from the vector between the midpoint **m** of **x_skewed_** to the midpoint between the wing hinges. The plane of reflection was defined by its normal **n_r_**, which was derived from the cross product between the y-axis vector y and the x-axis of camera coordinate system **c**, i.e. the posterior-anterior axis of the flies in global camera coordinates. To ensure orthogonality between the x- and y-axis, the vector y was redefined by calculating the cross product between vector **x** and the initial vector **y** based on the wing hinges. Eventually, the z-axis was derived from the cross product of vectors y and x to obtain a complete right-handed coordinate system.

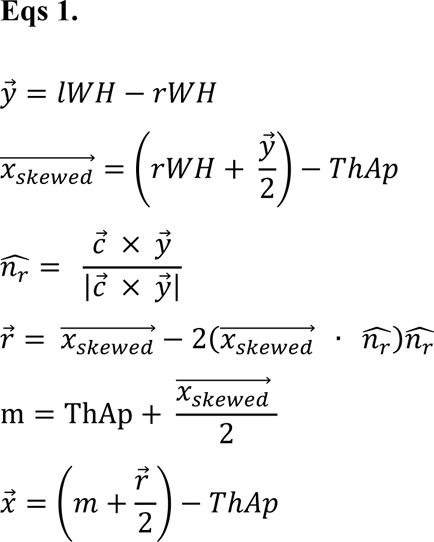

### 2.5 Detection of swing and stance phases

To compare model errors and joint movements between different flies, we normalized the time courses of the swing and stance phases. For this, we determined the lift-off and touchdown events for each step of all legs. Since the distance between tarsus tip and the center of the ball of the motion capture setup must be equal to the radius of the ball, i.e. 3 mm, during stance phase, we performed a threshold operation to evaluate in which phase each leg was for each video frame. The center position of the ball was obtained by using an optimization function which minimized the distance of all tracked tarsus tips of all legs to the radius of the ball. A penalty factor of 100 was multiplied to tarsus tip distances smaller than the radius in the cost function. That prevented that tarsus positions were located inside the calculated ball. The threshold was set manually for each leg of each fly to ensure correct determination of lift-off and touchdown events. For data normalization with equal durations ranging from zero to one for step phases, data sets for each swing and stance phase were linearly interpolated to a sample size of 100.

### 2.6 Kinematic leg model based on anatomical landmarks

To model the motion of all six legs, we defined each leg as a kinematic chain consisting of multiple joints connected by rigid segments of a specified length. Each kinematic chain comprised the same joints which we tracked with our motion capture system, i.e. ThCx, CxTr, TrFe, FeTi, TiTar, and Tar (see 2.3). For this, joint condyles positions and leg segment lengths were extracted from a high-resolution synchrotron radiation µCT scan carried out at the Paul-Scherrer Institute (PSI, Villigen, Switzerland). A single adult female wild-type *Drosophila melanogaster* specimen was scanned at the TOMCAT beamline of the PSI in absorption contrast at 10 keV with 20x magnification. The fly, including the legs, was segmented as a surface model (**Figure 2A**) from the image stack using ITK-snap (v.3.6; Yushkevich et al., 2006) with one label for each leg segment and major body part. The resulting stl-file was exported to Blender (v.2.79b) to identify the centers of each joint condyle of each leg segment in 3D. The main rotational axis and center position of each joint were derived from the vector between the joint condyles and its center, respectively, while the leg segments were defined by the vector between consecutive joint positions or the Tar. Due to bilateral symmetry of the insect bauplan, the front, middle, and hind leg of the right body side were analyzed and positions were mirrored to the left side. Since the actual DOF configuration of the leg joints in *Drosophila* is not yet fully known, each model joint was first defined as ball-and-socket joint equipped with three DOFs, namely yaw, pitch, and roll (**Figure 2C**). The yaw axis was defined by the main rotational axis of the joint. The roll axis was specified by the leg segment vector controlled by the joint to allow longitudinal rotations, while the pitch axis was defined by the cross product of the yaw and roll axes to allow the full range of motion of a spherical joint. Importantly, each DOF could be set as mobile or fixed in our inverse kinematic solver, so that they either could contribute to leg movements or were arrested in a constant position. Additionally, the positions of the ThAp, lWH, and rWH were extracted from the µCT data and served to define a body coordinate system for the model.

**Figure 2.**
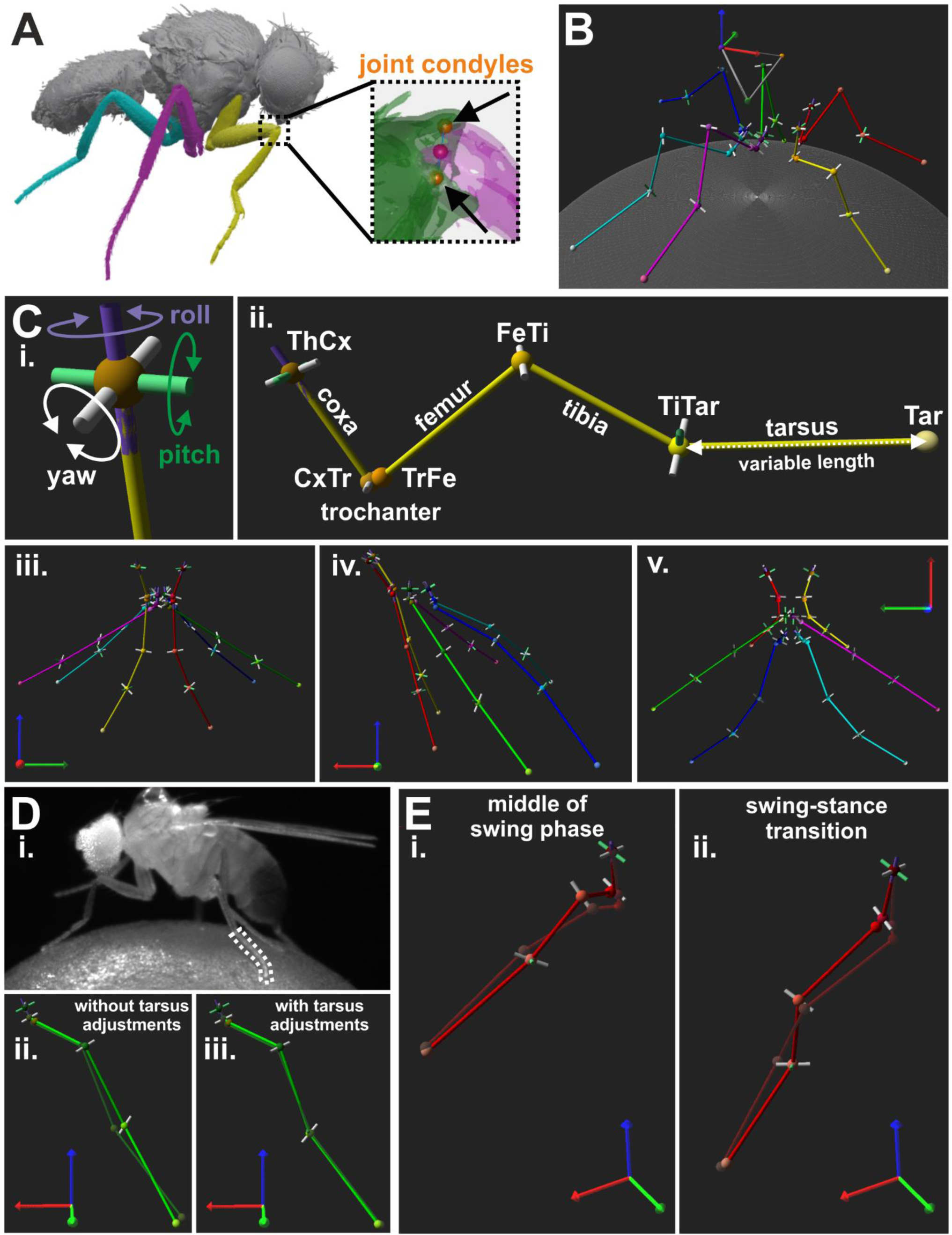
Creation of a kinematic reference leg model for *Drosophila*. (**A**) Image of µCT scan used to extract positions of joint condyles. Magnified view on the femur-tibia joint (FeTi) with highlighted joint condyles are displayed in the dashed box. (**B**) An exemplary walking posture of the reference leg model. (**C**) DOF configuration and initial posture of the leg model. **i.:** Directions of motion for a model joint with all three DOFs, i.e. yaw (white), pitch (green), and roll (purple). **ii.:** Kinematic leg chain of the right front leg of the reference model. **iii.-v.**: Initial posture of the model as seen from the front (**iii.**), side (**iv.**), and top (**v.**) view. (**D**) Example for bending of the tarsus during the stance phase and its influence on the model. **i.**: Video frame showing a walking fly with a bent middle leg tarsus. **ii.-iii.**: The resulting model leg postures without (**ii.**) and with adjustments for tarsus fitting (**iii.**), i.e. additional pitch DOF in the tibia-tarsus joint (TiTar) and adjustment of the tarsus lengths for each leg posture. Solid light green and transparent dark green legs chain represents the joint and segment positions for the model and the motion captured legs, respectively. (**E**) Two examples for front leg misalignment of the model’s tibia with its motion captured counterpart at the middle of the stance phase (**i.**) and the transition from swing to stance (**ii.**).

Since alignment of leg segments was predetermined by the leg postures in the µCT scan, we adjusted the joint and tarsus tip positions to obtain a better interpretable initial model posture (**Figure 2C**). For this, we adjusted the angles of the yaw DOFs in our kinematic chains until the forward kinematics (see 2.7) resulted in extended leg segments and stored the new positions of the joint condyles for construction of all subsequent models, i.e. the initial angles were zero in the kinematic chains for these initial leg postures. We did not perform additional adjustments for the angles of pitch and roll DOFs because this would have disrupted the anatomical relationship between leg segments and joint rotational axes.

To construct an orthogonalized model version with main rotational axes perpendicular to the leg segments, the original joint rotational axes were replaced by the body coordinate system axis that best represented the main rotational direction of the original yaw DOF (**supplemental video 5**). For this, new joint condyles positions were determined based on the joint center position ± the respective body coordinate system axis. Furthermore, the leg segments were linearized, i.e. pointing straight down from the ThCx, by displacing the joint positions based on the segment length in the original model.

### 2.7 Forward kinematics of the kinematic leg model

To solve the forward kinematics of the leg chains, i.e. determining the position and orientation of each joint and the tarsus tip by a given set of joint angles, all joint DOFs and the tarsus tip were represented by local coordinate systems (LCS) which were constructed according to the standard Denavit-Hartenberg (D-H) convention (Denavit and Hartenberg, 1955; Spong et al., 2020). For this, the z-axis of the LCS for each DOF was derived from its rotational axis, while the origin and x-axis depended on the relationship between the z-axis and the rotational axis of the previous DOF:

a. when both axes were not coplanar, the common normal, i.e. the line that orthogonally intersects both rotational axes, was used to specify the x-axis and the origin was the point of intersection on the z-axis.
b. when both axes intersected, the x-axis was defined by the normal of the plane formed by both axes and the origin was the point of intersection.
c. when both axes were parallel to each other, there were infinitely many common normals. In this case, the x-axis was derived from the common normal that passes through the joint center of the respective DOF which also served as origin.

The y-axis was derived from the cross product of the z-axis and x-axis to form a right-handed LCS. There were two LCS in the kinematic leg chain which could not be defined by this approach. Since there was no previous DOF for the first DOF in the kinematic chain, i.e. the ThCx, the x-axis was defined by the cross product of its z-axis and the x-axis of the body coordinate system and the ThCx position served as origin. Additionally, the tarsus tip was the end effector of the kinematic chain and thus it did not have a rotational axis. Here, the LCS of the TiTar-roll LCS was duplicated and its origin was translated to the tarsus tip position.

Afterwards, the four D-H parameters were calculated for the LCS’ of two consecutive DOFs to obtain a transformation matrix describing their geometrical relationship. The joint angle **θ** and the twist angle **α** were calculated by constructing two planes based on the axes of the LCS and determining the angle between their plane normal (**Eqs. 2**).

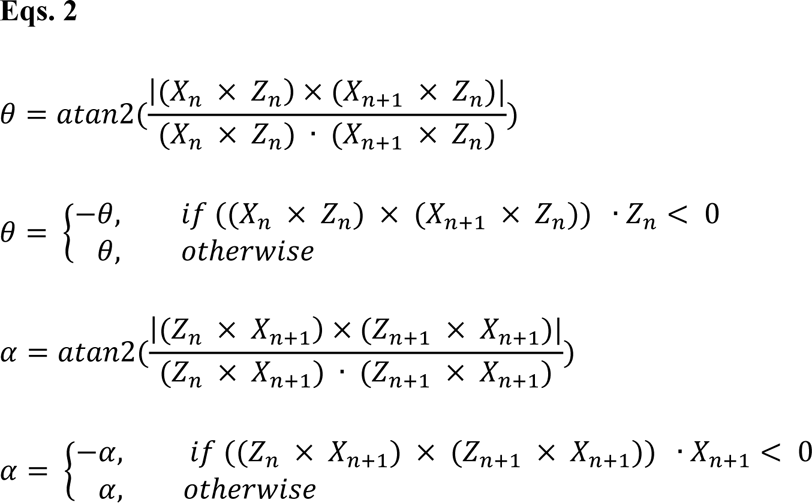

The link offset **d** and the link length **r** were derived from the intersection points **p_z_** of the common normal between the z-axes of the LCS or set to zero when both z-axes intersected (**Eqs. 3**).

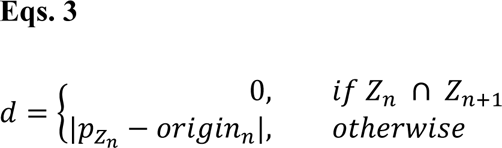

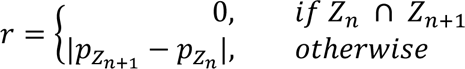

The D-H parameters were inserted in the transformation matrix **T** for each DOF. For calculating the forward kinematics for a given set of joint angles, the respective angle of a DOF was added to its initial **θ**. To obtain the final posture of a kinematic leg chain in global coordinates, the initial positions of the joint/tarsus tip was multiplied with the product of the transformation matrices from all preceding DOFs and the space coordinate transformation matrix **B_ThCx_** which was derived from the axis vectors and the origin of the LCS of the ThCx in global coordinates (**Eqs. 4**).

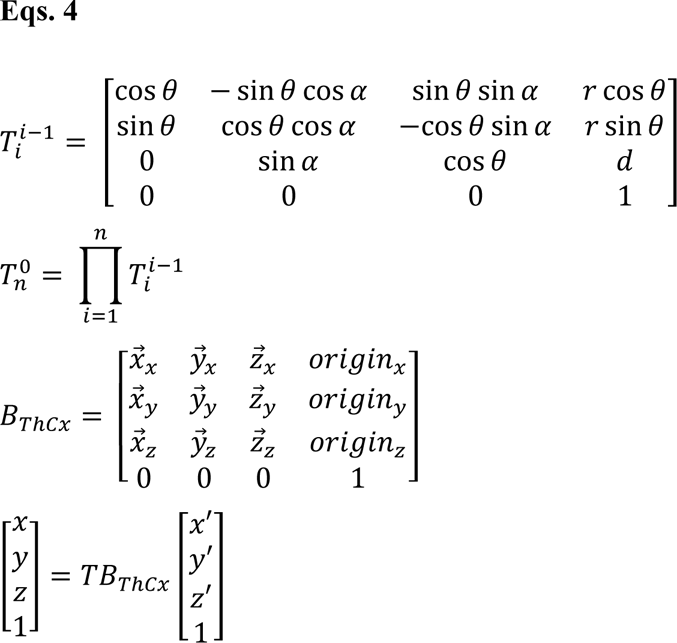

### 2.8 Inverse kinematics by global optimization

To determine the joint angle sets of leg postures exhibited by fruit flies during walking, the kinematic leg chains were fitted to the motion captured leg postures. For this, the forward kinematics of the leg chains were optimized by using an iterative conjugate gradient descent approach that minimized the sum of squared distances between the tracked and the modeled positions of the joints and the tarsus.

We applied a nonlinear trust-region reflective algorithm (Branch et al., 1999; from SciPy optimization package version 1.6.2) with a maximum iteration number of 100-times the number of mobile DOFs and 10^−3^ as termination criteria for changes in the cost function, resulting angles, and the norm of the gradient. Angle constraints for all DOFs (**supplemental table 1**) were introduced to prevent solutions that resulted in unnatural leg postures. Since the actual ROMs of joint DOFs in *Drosophila* are not known, angle constraints were empirically adjusted to ensure an appropriated model fit but minimizing solutions that reached the range limits. The cost function based on the distances between the CxTr, TrFe, FeTi, TiTar, and Tar positions of the model and their motion captured counterparts. To improve fitting, we weighted the distances of TrFe, FeTi, and TiTar positions more strongly than those of the other leg keypoints for the front (1.5, 2.0, 1.5) and middle (1.25, 1.5, 1.25) legs.

For each walking sequence, the size and leg segment lengths of the model were adjusted to the respective fly. For this, reference triangles were constructed based on the positions of the ThCx, rWH, and lWH of the model and the fly and a scale transformation matrix derived from these triangles was applied on the joint condyle positions of the model. The lengths of all leg segments of the model were then adjusted to the median length of their motion captured counterpart by linearly translating the joint condyle positions. Afterwards, the kinematic leg chains were constructed as described in **2.7**.

An initial set of angles was estimated to prevent that the global optimization process was directed to a sub-optimal local minimum solution. For this, the angles for each joint were individually optimized to fit the model joint position to the motion captured position in the first leg posture of a walking sequence. Afterwards, the optimization algorithm was applied globally, i.e. all model DOF angles were optimized simultaneously, to fit the model leg to the first leg posture. For all following leg postures of a walking sequence, the obtained angles from the previous leg posture served as initial estimate for the optimization process of the current leg posture.

### 2.9 Data analysis and statistics

All analysis routines, visualizations, and graphical user interfaces used were implemented in Python (3.9.5) or MATLAB (R2021a, The Mathworks, Natick, Massachusetts, USA). Data was expressed as mean ± standard deviation (SD), if not stated otherwise. While N denotes the number of experimental animals, n indicates the number of an analyzed feature. When the median was used, variability of the data set was indicated by the interquartile range (IQR) defined as the difference between the 75^th^ and 25^th^ percentiles.

To evaluate the model error, i.e. how well the kinematic leg model could adapt to the motion captured leg postures, we calculated the summed Euclidean distance between the CxTr, TrFe, FeTi, TiTar, and Tar positions of the model and their tracked counterparts. Since the joint and tarsus positions were in body coordinates at first, we converted them to global camera coordinates to obtain the model error in metric units, i.e. µm. We also calculated the area under curve (AUC) of the time courses of the model error as a measure for the goodness of fit between different DOF configurations. For this, we approximated the integral of the normalized time courses (see 2.5) of the mean model error during swing and stance phase by using Simpson’s rule.

Since we used the joint DOF angles obtained from our inverse kinematic solver, most of the angle values are reported in relation to the initial posture of the model (**Figure 2C**). To simplify the interpretation for movements of the two main leg joints, namely the CxTr and the FeTi, we post-processed the obtained angles of their yaw DOFs. Because the leg segments were fully extended in the initial leg postures of the model with an angle of 0° for these DOFs (see 2.6), we added 180° to the angles obtained from the solver. Therefore, a final angle of 0° or 180° described a fully flexed or fully extended posture, respectively. To combine time courses of joint angles for contralateral legs, the sign of angles for the yaw and roll DOFs of the left legs were inverted, but not for angles of pitch DOFs. The ROM of joint DOFs was derived from the maximum and minimum observed angle for each DOF. For analysis of rotations about the femur-tibia plane, we determined the plane normal by calculating the cross product between the vectors of the femur and the tibia. The ROM of femur-tibia plane rotations was the angle between the plane normals of the first and last leg posture of an analysed step cycle phase.

## 3 Results

### 3.1 Creation of a kinematic reference leg model for *Drosophila*

To test for the DOF requirements to accurately model the kinematics of *Drosophila* legs for straight walking, we first constructed an initial reference model based on reported DOFs for *Drosophila* and other insects (Cruse and Bartling, 1995; Soler et al., 2004; Bender et al., 2010; Goldsmith et al., 2022; Lobato-Rios et al., 2022) (**Figure 2B-C**). The ThCx was implemented as ball-and-socket-like joint with all three DOFs, i.e. yaw, pitch, and roll. Since the CxTr and FeTi are considered to be hinge joints in insects (Cruse and Bartling, 1995; Full and Ahn, 1995; Frantsevich and Wang, 2009), we only used the yaw DOF for those in our model. Two previously published models for *Drosophila* legs suggested that a roll DOF exists in either the TrFe (Goldsmith et al., 2022) or the CxTr (Lobato-Rios et al., 2022). However, to test if such an additional DOFs or joint was still necessary to model straight walking in our model with rotational axes based on anatomical landmarks, we chose to omit this DOF and initially fixed the whole TrFe and the CxTr-roll DOF. Because of their small size and proximity, motion capture of all individual tarsal joints was not feasible. We therefore had no guidance points for fitting a tarsus with five individually articulated segments and modelled the tarsus as a single segment. However, in initial modelling attempts we had difficulty to fit our model to the motion capture data when we modelled the tarsus with only an active yaw DOF in the TiTar and a constant length of the tarsus. This was mainly due to the fact that the tarsus is relatively flexible and can bend during the stance phase, resulting in apparent length changes (**Figure 2D**). Consequently, the model tarsus was commonly too short or too long for an optimal fit, which also affected the fit of all other model joints. To compensate for this the length of the tarsus was adjusted to the length of the tracked tarsus of the animals for each leg posture. We further added the pitch DOF to the TiTar to account for the observed bi-directionality of tarsus bending. In this way, we were able to model the complexity of tarsal motion with only three parameters.

Equipped with this basic model, we measured how well it could adapt to natural leg postures by applying inverse kinematics to fit the model to the recorded motion capture data of 12 flies (five females and seven males). The summed mean model error, i.e. the distance between the model and the tracked animal leg parts locations, was largest for the front legs (**Figure 3A**). Moreover, the mean model error was not consistent during the step phases, but showed a minimum of 342 ± 61 µm at the middle of the stance phase, increased steadily throughout the remaining stance phase and the swing phase to a maximum of 560 ± 80 µm, before it started to decrease at the onset of the stance phase (n, swing/stance: 232/213). The increase in error was largely due to the inability of the model to accurately replicate the positions of the TrFe, FeTi, and TiTar, which also resulted in misalignment between the model and the tracked leg segments. During the swing phase, this was particularly noticeable as the model’s tibia was not properly aligned with its tracked counterpart, causing the two to cross over each other (**Figure 2E, supplemental video 2**). This also adversely affected the model’s ability to reach the tracked tarsus tip positions, resulting in a 3.5-fold greater distance of the model tarsus tip at the end swing phase.

**Figure 3.**
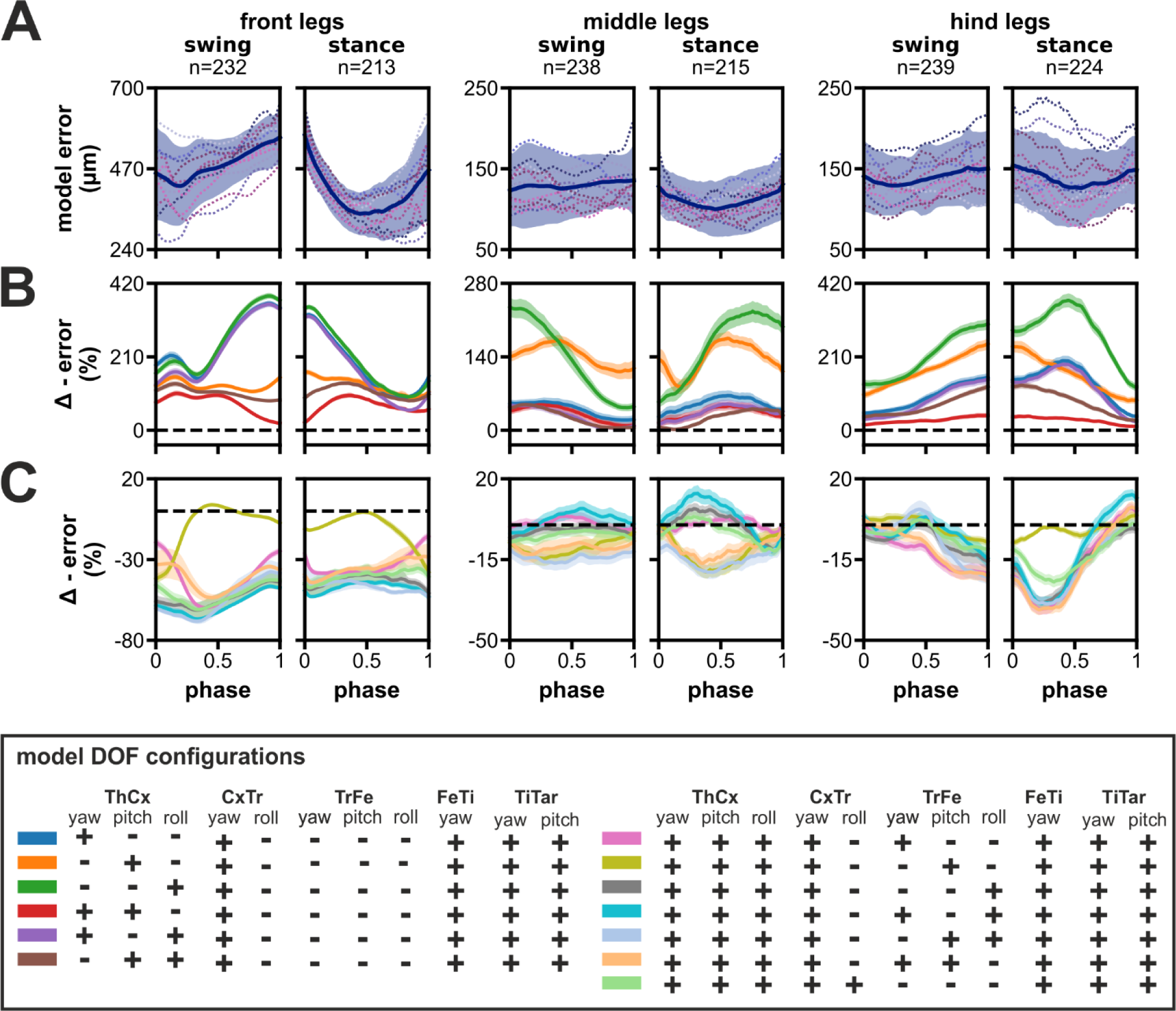
Model error and the impact of reducing the number of DOFs in the ThCx or adding DOFs to the TrFe and CxTr. (**A**) The time course of the summed mean error for front, middle, and hind legs during swing and stance phases. Blue line and area represent the mean ± SD, while dashed colored lines display the mean time course of individual flies (N=12). (**B-C**) Relative change of model error by removal of DOFs in ThCx (**B**) or addition of DOFs in the TrFe/CxTr (**C**) compared to the reference model. Colored lines and area display the mean change ± 95% CI. Dashed black lines indicate zero change, i.e. no difference to the error of the reference model.

In contrast, the summed mean model error for the middle and hind legs was not only substantially smaller, but also relatively constant during both step phases (**Figure 3A**). The error was similar for middle and hind legs and ranged from 100 ± 34 µm to 136 ± 46 µm (n, swing/stance: 238/215) and 126 ± 47 µm to 154 ± 49 µm (n, swing/stance: 239/224), respectively. Additionally, a proper alignment of model and tracked leg segments was achieved throughout the step cycle (**supplemental video 2**). Taken together, the reference model was already capable of adopting the tracked leg postures of the middle and hind legs exceedingly well, but was not able to sufficiently match leg postures of the front legs.

### 3.2 Impact of reducing the number of DOFs in the ThCx or adding DOFs to the TrFe and CxTr on the model error

Although the ThCx is assumed to have three DOFs in *Drosophila* legs (Lobato-Rios et al., 2022), it was reported that insects might mainly use only one or two DOFs for straight walking, particularly in the middle and hind legs (Hughes, 1952; Cruse and Bartling, 1995; Cruse et al., 2009; Bender et al., 2010). Therefore, we investigated whether our model required all three DOFs in the ThCx to adequately reproduce natural leg stepping postures. For this, we evaluated the difference in model error between the reference model and other DOF configurations with fewer DOFs in the ThCx. The elimination of DOFs in the ThCx generally resulted in a larger model error for all three leg pairs with a 1.2 to 3.3-fold larger AUC of the absolute model error time course (**supplemental table 2**). When we analyzed the relative change of the summed mean model error times courses, we found that individual DOF configurations performed differently in each leg pair throughout the step cycle (**Figure 3B**).

For the front legs, fixing the ThCx-pitch DOF led to the largest increase in model error (57% to 384%) which peaked for all affected DOF configurations at the transition between swing and stance phases. When only the pitch DOF was mobile in the ThCx, the increase in error was less pronounced but still substantial (94% to 167%); addition of either the yaw or roll DOF did not considerably reduce the increase in error over most of the step cycle. Interestingly, the relative increase in error was only 11% at the stance-to-swing transition for a model configuration with mobile yaw and pitch DOFs, suggesting that these DOFs were mainly required for modelling postures of the front legs at this stage.

For the middle legs, however, the ThCx-yaw DOF was more important for model fitting. The model error was similar for ThCx configurations with a moveable yaw DOF which was 4% to 66% higher than the reference leg model error. In contrast, the model error was largest for ThCx configuration with only the pitch or roll DOF, resulting in an increase in error of 83% to 175% or 43% to 235%, respectively. However, when the pitch and roll DOFs were moveable, while the yaw DOF was absent, the increase in model error was slightly smaller with 1% to 49% compared to model configurations employing the ThCx-yaw DOF.

For the hind legs, fixing the roll DOF had the smallest effect on the error, i.e. 11% to 43% larger error, whereas the increase in error was greatest, i.e. we observed a 124% to 371% larger error, when the roll DOF was the only moveable joint DOF. Although the model error steadily increased during the swing phase for all model configurations, the increase in error peaked at different time points when the pitch DOF was moveable or fixed. While model configurations with a movable pitch DOF showed the largest increase in error at the transition from swing-to-stance phase, model configurations with a fixed pitch DOF showed a transient increase in the relative error time course at the middle of the stance phase. This suggests that the pitch DOF is important for modelling postures of the hind legs at this stage of the step cycle.

Next, we examined if a moveable TrFe could improve the model performance (**Figure 3C**). In addition, we also tested a DOF configuration in which the CxTr had a roll DOF in addition to its yaw DOF as this configuration was suggested for *Drosophila* by Lobato-Rios et al. (2022). For the front legs, addition of an extra DOF resulted in substantial decrease in model error throughout the step cycle with a 1.6 to 2.2-fold smaller AUC of the absolute model error time course (**supplemental table 2**), except when only the pitch DOF was added to the TrFe. The largest reduction in model error was achieved when the TrFe or the CxTr were equipped with a movable roll DOF (relative change in error: TrFe with roll DOF: −38% to −68%; CxTr with roll DOF: −35% to −61%). Strikingly, the addition of a roll DOF also eliminated the misalignment between the model and the tracked tibia as observed in the reference model, which was not the case with the other DOF configurations that lacked such a roll component. Although we also observed a reduction in error by the addition of an extra DOF in the middle and hind legs, the amount of relative change was smaller compared to the front legs.

For the middle and hind legs, although we observed a reduction of the AUC of the absolute model error time course by up to 18% or 19%, respectively, for some DOF configurations this reduction might be rather negligible considering the already relatively small model error of the reference model for these leg pairs (**Figure 3A**). Furthermore, many DOF configurations did not consistently improve the model fit throughout the step cycle (**Figure 3C**). For the hind legs, however, almost all configurations with an additional DOF exhibited a reduction in error of more than 15% over a longer period during the step cycle; the largest reduction was observed in the first half of the stance phase. In contrast, a similar amount of reduction was only found for the middle legs when the TrFe had a movable pitch and roll DOF. Interestingly, while addition of a pitch DOF had the smallest effect on the model error for the front and hind legs, all DOF configurations equipped with a pitch DOF showed the largest reduction in error with up to 21% for the middle legs.

Although these findings suggested that elimination of some DOFs in the ThCx increases the model error only slightly, at least for the middle and hind legs, and that addition of an extra DOF in the TrFe can improve model fitting, it was not possible to deduce a unique DOF configuration for all leg pairs based on our data. Since we already obtained an appropriate model fit for the middle and hind legs without a mobile TrFe joint, we decided to fix this joint in our model in these legs for further analysis. In contrast, for the front legs an additional DOF was necessary to achieve an adequate model fit. We therefore added a mobile roll DOF in the TrFe because this configuration not only improved model performance the most with the least number of DOFs, but also resulted in a correct alignment between the model and the tracked tibia (**supplemental video 3**).

### 3.3 Angle time courses and movement directions of joint DOFs in the kinematic model

To examine the joint kinematics of our model in more detail, we analysed the angle time courses of each DOF and their relevance for the motion of the associated leg segment (**Figure 4**). However, we omitted the TiTar joint in our analysis because our modelling approach for tarsal movements allows only limited conclusions about the natural contribution of TiTar motion to leg stepping. To specify movement directions, we used the following anatomical terms of motion (Hughes 1952): for the coxa, promotion and remotion describe the anterior and posterior movement in relation to the body, while adduction and abduction refer to lateral movement either towards or away from the body midline. Rotations of segments are specified by the main anatomical direction in which the following segments were moved, i.e. medial or lateral rotations move the subsequent segments towards or away from the body midline, while anterior or posterior rotations moved the subsequent segments towards the head or abdomen of the animal, respectively. The relationship between two adjacent leg segments was described by flexion and extension which results in a decrease or increase, respectively, of the inner angle between the segments. Additionally, leg levation and depression describes movements in which the leg is lifted off the ground or lowered to the ground.

**Figure 4.**
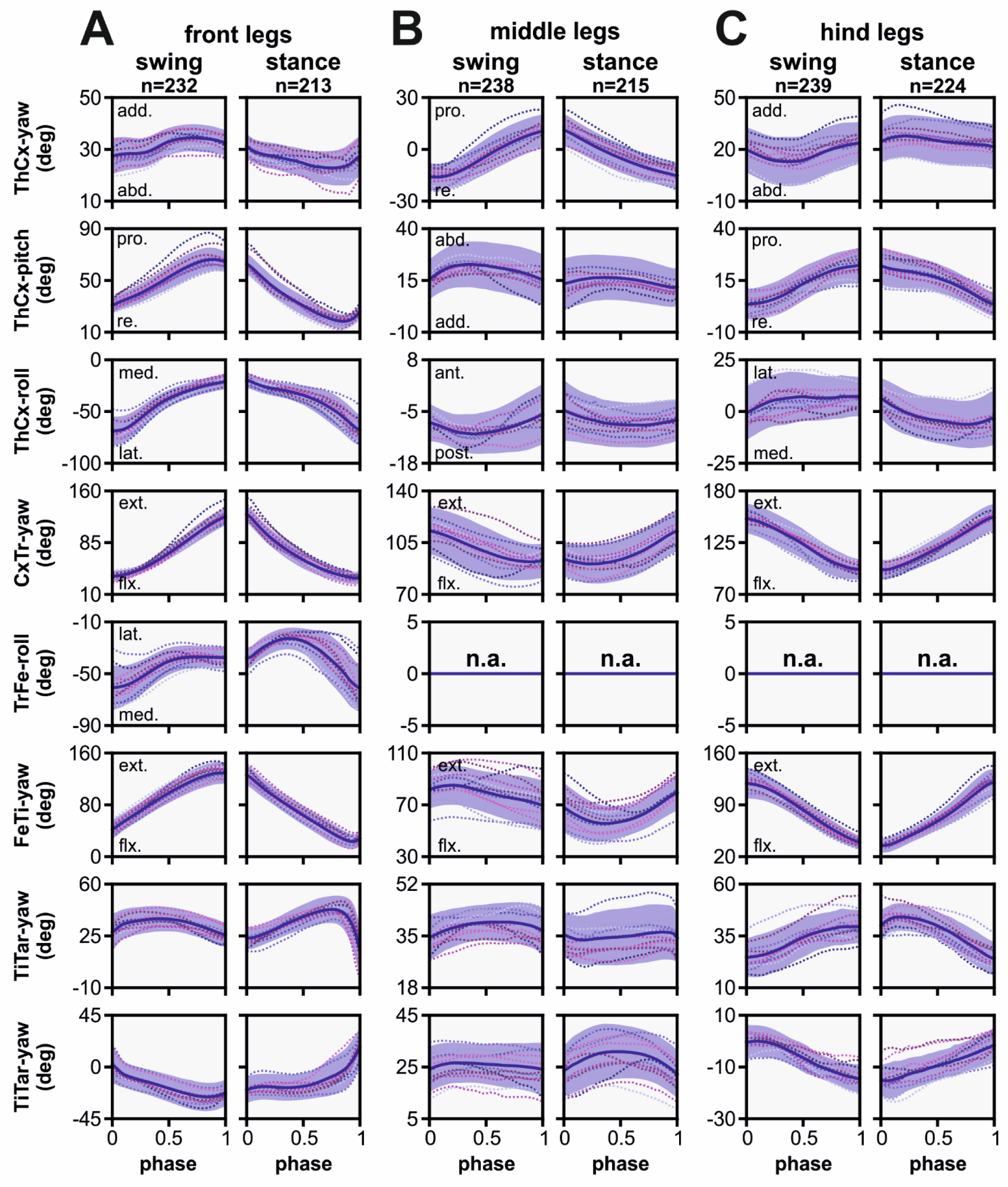
Angle time courses and movement directions of joint DOFs in the kinematic model. The angle time courses are shown in relation to the normalized swing and stance phase for front (**A**), middle (**B**), and hind (**C**) legs. Angles were calculated in relation to the initial posture of the model, except for CxTr-yaw and FeTI-yaw DOFs. These were post-processed to show the relationship between the linked segments: 0° indicates a complete overlap of both segments, i.e. maximally flexed, while 180° indicates that both segments are co-linear, i.e. maximally extended. Blue line and area represent the mean ± SD, while dashed colored lines display the mean time course of individual flies (N=12). Abbreviations: abd., abduction; add., adduction; pro., promotion; re., remotion; ext., extension; flx, flexion; med., medial rotation; lat., lateral rotation; ant., anterior rotation; post., posterior rotation.

Generally, the movement patterns of forward stepping showed distinct leg pair-specific kinematics. Forward stepping of the front and hind legs was primarily executed in the anterior-posterior plane, i.e. the legs were mainly extended or flexed by combined actions of the ThCx, CxTr, and FeTi in addition to leg levation and depression. However, the respective leg movements occurred in opposite step cycle phases. For instance, extension and contraction of the leg was performed during swing and stance phase, respectively, in the front legs. In contrast, in the hind legs, contraction of the leg was performed in the swing phase, while extension of the leg was observed in the stance phase. The middle legs, on the other hand, showed an idiosyncratic movement pattern. Here, protraction and retraction of the leg were achieved by promotion and remotion of the coxa as well as a rotation of the femur-tibia plane that resulted in an anteriorly or posteriorly pointing of the tarsus tip at the end of the swing or stance phase, respectively (see also 3.4.). Additionally, levation and depression of the leg was mainly governed by the CxTr and flexion and extension of the tibia was less pronounced compared to the other leg pairs.

In the front and hind legs, promotion and remotion of the coxa was mainly driven by the ThCx-pitch DOF, while the ThCx-yaw DOF performed that movement in the middle legs. As found in other insects, promotion and remotion occurred in all leg pairs during the swing and stance phase, respectively. Although the shape of angle time courses was similar, the ROM observed for the individual leg pairs differed with 40.87° ± 5.21°, 26.48° ± 4.58°, and 18.60° ± 3.85° for the front, middle, and hind legs, respectively. In contrast, adduction and abduction was governed by the yaw DOF in the front and hind legs and by the pitch DOF in the middle legs. While the ROM was comparable for all leg pairs (front legs: 8.24° ± 2.74°, middle legs: 7.63° ± 2.85°, hind legs: 8.85° ± 2.98°), we found some variations in the course of adduction and abduction between the leg pairs. For the front legs, adduction occurred relatively rapidly at the middle of the swing phase and at the end of the stance phase, whereas gradual abduction was mainly observable from the end of the swing phase to the end of stance phase. In contrast, the hind legs exhibited adduction from the middle of the swing phase to the first quarter of the stance phase, followed by gradual abduction to the end of the stance phase and faster abduction in the first half of the swing phase. Interestingly, a complete cycle of adduction and abduction occurred in each of swing and stance phases for the middle legs, but was more pronounced in the swing phase than in the stance phase. Although the ThCx-roll DOF was used by all three leg pairs, it was most extensively used by the front legs (ROM: 40.87° ± 5.21°) and only to a lesser amount by the hind (ROM: 11.27° ± 3.01°) and middle (ROM: 4.91° ± 2.67°) legs. Additionally, while the ThCx-roll DOF governed medial and lateral rotations for the front and the hind legs, it drove posterior and anterior rotations in the middle legs.

In the front legs, the additional TrFe-roll DOF was used for lateral rotation of the femur-tibia plane during the first half of the swing phase. This lateral rotation was continued during the first half of the stance phase before it switched to medial rotation for the remaining stance phase. The CxTr-yaw and the FeTi-yaw DOFs were used for flexion and extension of the trochanter and the tibia, respectively. These movements were generally more pronounced in the front and hind legs than in the middle legs (ROM of CxTr/FeTi for front, middle, and hind legs: 91.24° ± 9.54°/ 96.55° ± 7.65°, 22.51° ± 3.38°/ 21.52° ± 5.43°, 56.34° ± 9.00°/ 84.11° ± 9.94°). Furthermore, while the front legs showed extension and flexion during the swing and stance phase, respectively, these movements were performed in the opposite phase for the hind legs. For the middle legs, however, flexion and extension of the trochanter occurred in swing and stance phase, but flexion of the tibia was observable almost during the entire swing phase and continued in the first half of the stance phase, while extension of the tibia occurred mainly in the second half of the stance phase. Additionally, flexion of the tibia in the middle legs was not gradual as compared to the front and hind legs, but it was more prominent at the early stance phase.

### 3.4 Rotations of the femur-tibia plane in the middle legs were driven by two yaw DOFs

Remarkably, although femur-tibia plane of the middle legs in *Drosophila* showed pronounced rotational movements during forward walking (Karashchuk et al. 2021; **supplemental video 4**), our model did not require any additional roll DOF in the TrFe for the adoption of the postures resulting from that rotational movement, unlike the front legs. To identify the kinematic origin of these femur-tibia plane rotations, we performed an *in silico* experiment in which we tested the ability of single as well as combinations of DOFs proximal to the femur to rotate the plane (**Figure 5**). For this, we set the middle legs of the model to their initial posture of each analysed swing (n=238) or stance phase (n=215) by using all available DOF angles and subsequently updated only the angles of selected DOFs for the remaining leg postures of the respective phase.

**Figure 5.**
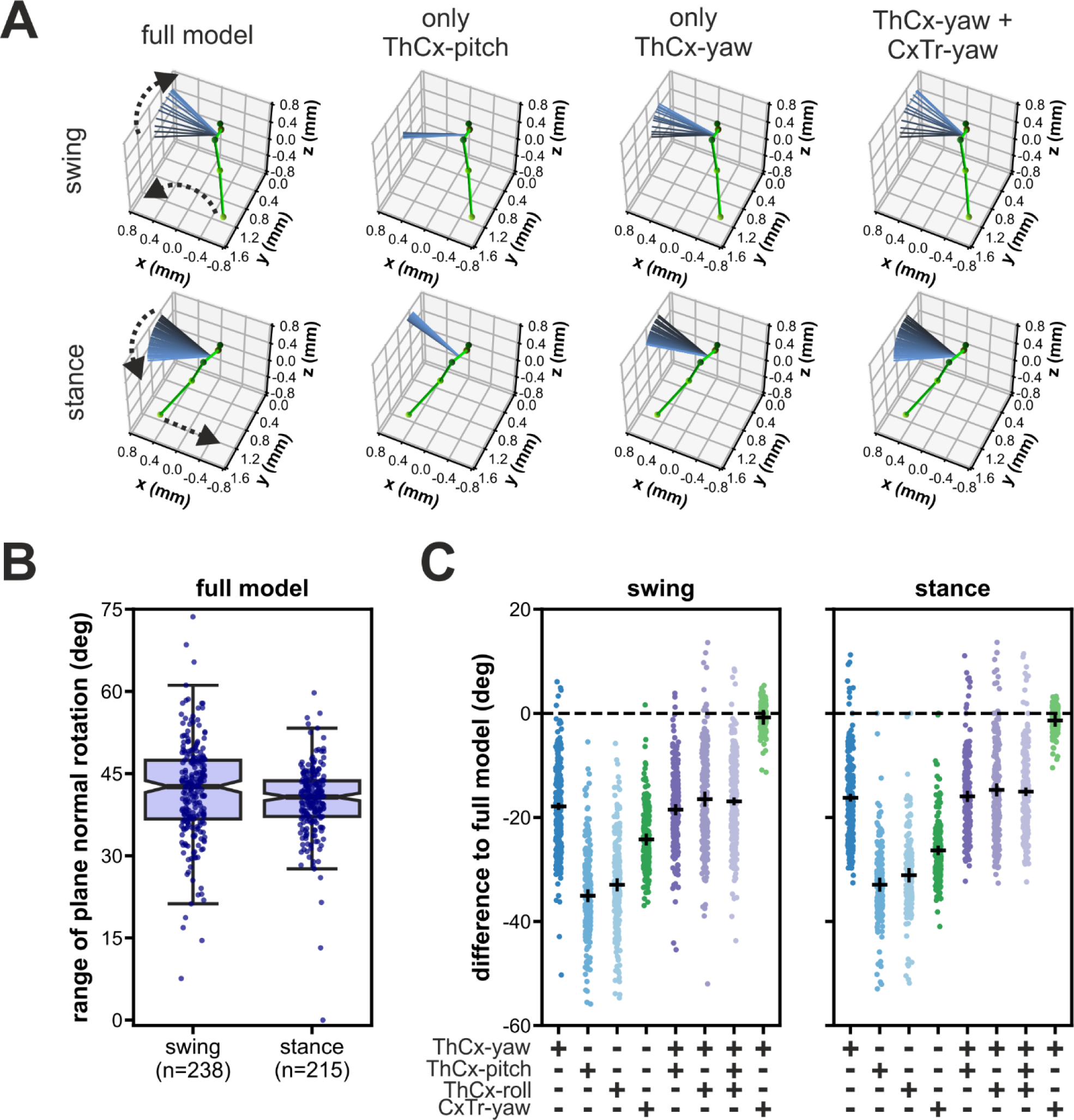
Rotations of the femur-tibia plane in the middle legs were driven by two yaw-DOFs. (**A**) Representative examples of a modelled left middle leg showing the effects on the femur-tibia plane rotations when only different combinations of joint DOFs proximal to the femur were updated after the model leg posture was initiated with all DOFs for a swing phase (upper panels) and a stance phase (lower panels). The initial leg posture is represented as ball-and-stick model in green. The orientation of the femur-tibia plane normal for all leg postures during the respective phase is displayed as colored-coded lines from black (initial posture) to light blue (final posture). Dashed arrows indicate the direction of the tarsus tip and the plane normal rotations during the respective phase. (**B**) Range of rotation of the femur-tibia plane normal in the full model, i.e. all DOFs were updated. (**C**) Absolute differences to the rotational range of the full model for all tested DOF configurations. Black solid lines represent the mean ± 95 CI.

When we updated all present model DOFs, i.e. we used the full established model, the median rotational range of the femur-tibia plane was 42.6° (IQR: 10.7°) and 40.7° (IQR: 6.5°) for the swing and stance phase, respectively (**Figure 5B**). Interestingly, almost all other tested DOF configurations resulted in a largely reduced rotational range (**Figure 5A+C**). We observed the largest reduction when only the pitch or roll DOF of the ThCx were updated (mean difference ± 95% CI for swing / stance: pitch DOF only: −35.1° ± 0.9° / −32.9° ± 1.0°; roll DOF only: −33.0° ± 0.9° / −31.1° ± 1.1°). In contrast, the rotational range was larger when the ThCx-yaw was the only moveable DOF, but it was still reduced by 17.9° ± 0.4° and 16.2° ± 0.4° for the swing and stance phase, respectively. Updating either the ThCx-pitch, or the ThCx-roll, or both DOFs in addition to the ThCx-yaw DOF did not enhance the rotational range (**Figure 5C**). Additionally, the reduction of the rotational range was 24.2° ± 0.8° for swing phases and 26.3° ± 0.6° for stance phases when only the CxTr-yaw DOF was updated in the model. Strikingly, combining the ThCx-yaw and the CxTr-yaw DOFs resulted in an almost complete recovery of the rotational range (mean differences ± 95% CI for swing / stance: −0.8° ± 1.1° / −1.4° ± 0.9°) (**Figure 5C**), showing that these two DOFs were the main contributors to the rotation of the femur-tibia plane of the middle legs in our kinematic model.

### 3.5 Effects of oblique joint rotational axes based on anatomical landmarks on kinematic modeling

Since a unique feature of our kinematic leg model was that yaw DOF rotational axes were derived from the positions of the joint condyles, we asked how their anatomical orientation improved fitting to the motion captured leg movements. For this, we selected three different DOF configurations of our model: the reference configuration, one with an additional TrFe-roll DOF, and one with an additional CxTr-roll DOF. We then compared these to alternative versions in which the rotational axes were orientated orthogonally to the leg segments (**supplemental video 5)**. We found that the orientation of most yaw DOF rotational axes of our model deviated, sometimes substantially, from their orthogonalized version. This consequently affected the direction of their rotational motion (**Figure 6A**).

**Figure 6.**
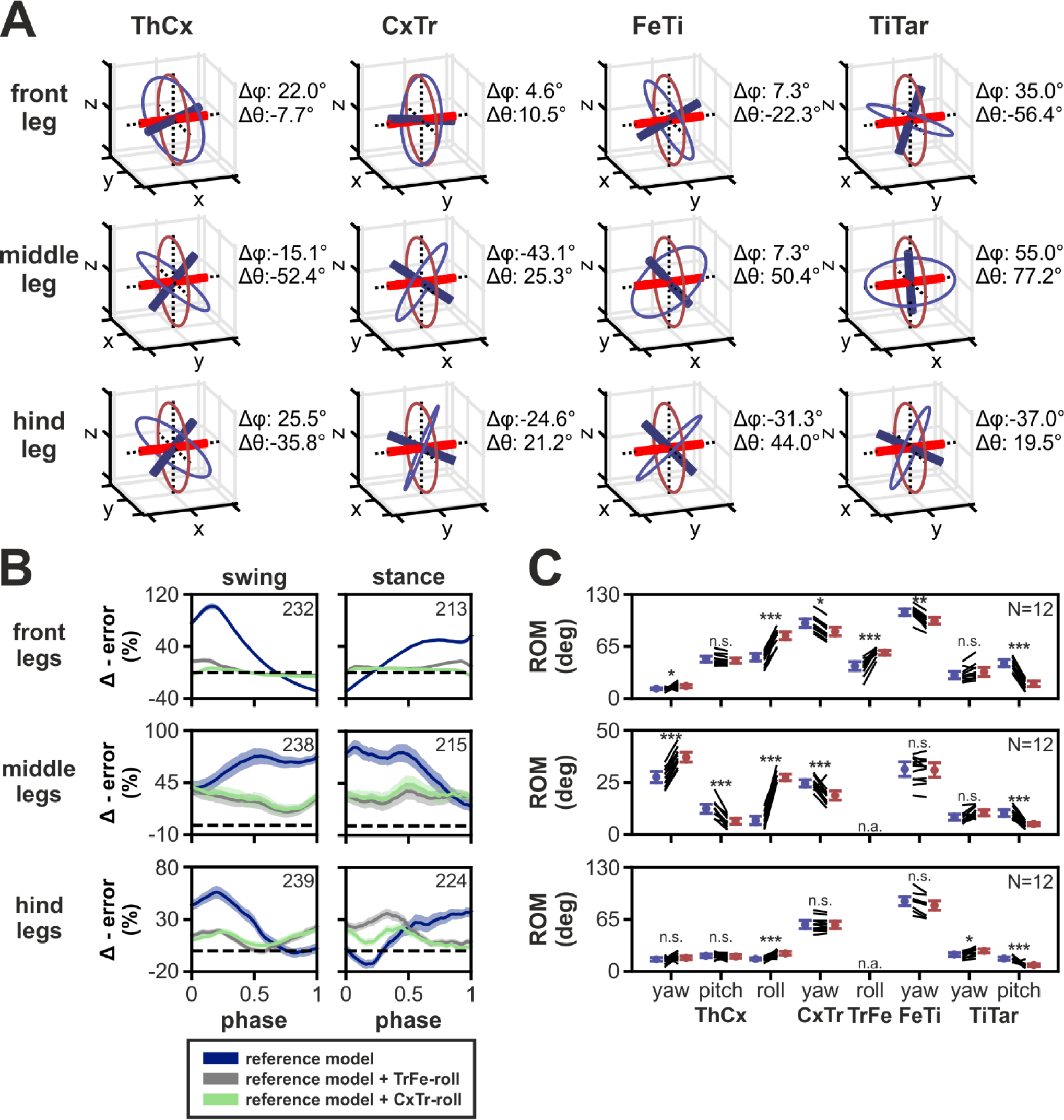
Effects of using oblique joint rotational axes based on anatomical landmarks on kinematic modeling. (**A**) 3D representations of the orientation for rotational axes based on positions of joint condyles (blue) compared to the model with axes orientated orthogonally to the leg segments (red). Colored circles represent the rotational motion the axes. Note that TrFe was omitted because only the TrFe-roll DOF axis was used in our final model which is the trochanter in both models. The differences in orientation between the axes are indicated by Δϕ and Δθ which based on azimuthal and inclination angles from the polar coordinates of the axes, respectively. (**B**) Relative change of model error of three different DOF configurations of the orthogonalized model compared to the model based on joint condyles positions for all three leg pairs. Colored lines and area display the mean change ± 95% CI and the same color code is used as in Figure 3. Numbers of analyzed step phases are indicated as numbers in the boxes. Dashed black lines indicate zero change, i.e. no difference in error. (**C**) Comparison of range of motion (ROM) of joint DOFs from our final model with a TrFe-roll DOF (blue) compared to the orthogonalized model version (red) for all three leg pairs. Colored lines represent the mean ± 95% CI, while black lines show the change in ROM for individual animals (N=12). Independent t-test results are displayed as: n.s. (not significant): p > 0.05; *: p < 0.05; **: p < 0.01; ***: p < 0.001. Abbreviation: n.a., not available.

For models with the reference DOF configuration, the orthogonalized model showed a larger error for all three leg pairs during almost the complete step cycle (**Figure 6B**). The maximum increase in relative mean error was 102%, 83%, and 56% for the front, middle, and hind legs, respectively. However, the orthogonalized model version performed better for the front and hind legs at the transition between swing and stance. For the front legs, the relative error decreased up to 29% at the end of the swing phase before increasing again at the begin of the stance phase, while for the hind legs the relative error was similar to our model at the end of the stance phase, but decreased up to 13% at the begin of the stance phase. Additionally, the orthogonalized versions of models with either a TrFe-roll or CxTr-roll DOF in the kinematic leg chains also displayed a larger model error compared to our model with these additional DOFs which was, however, less pronounced than we found for the reference DOF configuration (maximum increase in relative mean error for front, middle, and hind legs: with TrFe-roll: 19%, 40%, and 35%; with CxTr-roll: 7%, 44%, and 22%).

We further found differences in the ROM of several joint DOFs between our final model with an additional TrFe-roll DOF in the front legs and the respective orthogonalized model (**Figure 6C**). Although we observed reductions in the ROM for DOFs in the orthogonalized model in the front legs (mean differences ± 95% CI: −10.3° ± 0.7°, −10.7° ± 2.3°, and −25.8° ± 3.4° for CxTr-yaw, FeTi-yaw, and TiTar-pitch DOFs) and the middle legs (mean differences ± 95% CI: −6.1° ± 0.15°, −5.8° ± 2.5°, and −5.2° ± 1.3° for ThCx-pitch, CxTr-yaw and TiTar-pitch DOFs), we observed larger increases of the ROM in the available roll DOF in both leg pairs which was 26.8° ± 3.7° and 16.7° ± 3.9 for ThCx-roll and TrFe-roll DOFs in the front legs and 20.6° ± 1.4° for the ThCx-roll DOF in the middle legs. For the hind legs, changes in ROM were subtler in the hind legs, but the ROM for the ThCx-roll DOF was also increased by 7.1° ± 2.7°. Additionally, the ROM of the ThCx-yaw DOF in the middle legs showed also an increase of 9.5° ± 1.6°.

## 4 Discussion

In the present study, we introduced a kinematic 3D leg model for *Drosophila* with oblique joint rotational axes derived from anatomical landmarks. Our model was able to adopt natural leg postures occurring during forward stepping with high accuracy and, at the same time, revealed differences in the joint kinematics between the leg pairs. Thereby, our model further provides insights into biomechanical aspects of walking in *Drosophila*.

To the best of our knowledge, our model is the first 3D kinematic leg model of *Drosophila* in which the alignment of the main joint rotational axes was informed by the position of the joint condyles. The resulting rotational axes of the yaw DOF in all joints were therefore slanted and not aligned in an orthogonal orientation with regard to the leg segments (**Figure 6A**). By comparing our model with such oblique joint axes to a version with orthogonalized axes, which is a widespread modeling approach for kinematic leg chains in insects (Zakotnik et al., 2004; Petrou and Webb, 2012; Theunissen and Dürr, 2013; Goldsmith et al., 2022; Lobato-Rios et al., 2022), we found that our model had a smaller error in all three leg pairs most of the time. Additionally, the ROM for many DOFs, particularly for roll DOFs, differed substantially. One reason for the smaller required ROM of roll DOFs in our model might be that rotations around this type of DOF have a direct influence on the orientation of all subsequent rotational axes in the kinematic leg chain, suggesting that our model had to use these DOFs to a lesser extent to adapt the motion captured leg postures due to its more natural alignment of rotational axes. These findings also show that the resulting kinematics, such as joint angles, are strongly affected by the orientation of the rotational axes of the used DOFs, even when a kinematic model reproduces natural leg postures very well. This has further critical implications for the development of more sophisticated models of leg movements in *Drosophila*, such as dynamic, musculoskeletal, and full neuromechanical models, as deviations from the naturally occurring joint DOFs and their rotational axes could strongly affect the results (Stagni et al., 2000; Smith et al., 2012; Fohanno et al., 2013; Zuppke et al., 2023).

In our model, however, we could only align the yaw DOFs based on anatomical landmarks and used simplified assumptions to model pitch and roll DOFs. This means that the axes for roll DOFs were placed along the associated leg segment and axes for pitch DOFs were orientated orthogonally to both other DOF axes. However, these rotational axes might also be skewed in the fruit fly and thereby affect the model error and the range of motion used in our model. This notion is supported by the finding that the slanted and therefore non-orthogonal axes orientation between the two hinge joints that allow movements of the antennae in stick insects improves the efficiency of active tactile sensing (Krause and Dürr, 2004). For instance, our model achieved the best fit when all three DOFs were moveable in the ThCx for all leg pairs. Although the front legs in *Drosophila* likely use all three DOFs, as the coxa showed pronounced movements including extensive lateral rotations which is comparable to findings for the front legs of the cockroach (Kram et al., 1997; Tryba and Ritzmann, 2000; Ritzmann et al., 2004), many insects use only two or one DOFs of the ThCx in the middle and hind legs during forward walking (Hughes, 1952; Cruse and Bartling, 1995; Bender et al., 2010). It is therefore desirable to conduct future morphological and biomechanical studies to reveal the anatomical rotational axes of the other DOFs in joints with more than one DOF in *Drosophila*.

Another limitation of our model is the simplification of tarsal movements. The tarsus in *Drosophila* consists of five segments linked by ball-and-socket joints (Tajiri et al., 2010). Since it was not possible to track all tarsal segments with the motion capture system used here, we modeled the tarsus with two DOFs in the TiTar and adjusted its length for each leg posture, i.e. we effectively reduced 12 to 15 DOFs to three parameters. However, from a biomechanical perspective, the tarsus comprises a large percentage of the total leg length, serves as attachment structure, and has a passive spring effect which contributes to ground force transmission (Manoonpong et al., 2021). This suggests that a more detailed kinematic model of the tarsus is needed, for instance for dynamic simulations. On the other hand, tarsal segments do not have muscles that allow independent control by the nervous system, but are moved together by muscle tension on the long tendon (Soler et al., 2004). Consequently, our model with a simplified tarsus incorporates already the majority of joints which are under active control by the nervous system.

Although the TrFe is moveable in many insects (Frantsevich and Wang, 2009), its functional role in *Drosophila* is debated controversially, with opinions ranging from it being fused and immobile (Sink, 2006; Lobato-Rios et al., 2022) to at least limited mobility (Feng et al., 2020). We found that a roll DOF in the TrFe drastically improved fitting of our model to leg postures of the front legs, while it had much smaller impact on fitting of the middle and hind legs. This is in line with a kinematic model for crickets in which a mobile TrFe was required to model walking in the front legs, but not in the other leg pairs (Petrou and Webb, 2012). However, we observed the same improvement when we used a roll DOF in the CxTr as proposed by Lobato-Rios et al. (2022). Nevertheless, we favoured a mobile TrFe in our model, because the CxTr has commonly been characterized as hinge joint with only one DOF for leg levation/depression in insects (Cruse and Bartling, 1995; Kram et al., 1997; Tryba and Ritzmann, 2000; Büschges et al., 2008; Cruse et al., 2009) and the trochanter has muscles that are innervated by the nervous system in *Drosophila* (Soler et al., 2004; Enriquez et al., 2015). Their presence in this segment suggests at least some functional relevance in the context of leg movements.

Interestingly, the front and middle legs exhibited a rotation of the femur-tibia plane during forward stepping. While this rotation was very prominent in the middle legs and contributed to protraction and retraction of the leg, it was subtler in the front legs, though still necessary for adequate modeling of the leg kinematics. Femur-tibia plane rotations were previously reported for the middle legs in the cockroach (Bender et al., 2010) and *Drosophila* (Karashchuk et al., 2021). In contrast to the cockroach in which this rotation is proposed to emerge from actions of the TrFe (Bender et al., 2010), we found such a putative role for the TrFe only for the front legs in our model. In the middle legs, however, these rotations were exerted by the combined actions of the yaw DOFs of the ThCx and the CxTr, which can be most likely attributed to the alignment of their rotational axes.

One important finding from our model is that each leg pair showed distinct kinematics in terms of joint angles, ROM, and the necessity of individual DOFs for accurate fit of specific postures at different times throughout the step cycle. These observations have some implications for the demands of the underlying motor control system in *Drosophila* in terms of required muscle activation patterns and sensory feedback from proprioceptors. For instance, it is well-established that the femoral chordotonal organ (fCO) in insects encodes position, velocity, acceleration, and vibration of tibial movements (Hofmann and Koch, 1985; Hofmann et al., 1985; Matheson, 1990; Büschges, 1994; Stein and Sauer, 1999; Mamiya et al., 2018). However, although we found that the extent of angular changes of the FeTi was similar, tibial flexion occurred during the stance or swing phase for the front and hind legs, respectively, and vice versa during the opposite step phase. Additionally, tibial flexion and extension was much less pronounced in the middle legs. This suggests that the sensory signals originating in fCOs are leg-specific and this specificity must be accounted for when these signals are processed by the nervous system. This notion is supported by the findings of Chockley et al. (2022) who showed that inhibition of the fCO during walking in *Drosophila* strongly elongated the stance phase and trajectories of the front legs and the swing phase of the hind legs.

Another open question is to what extent the neuronal circuits underlying walking are conserved in insects. Since the nervous system and the morphology of legs evolved together, this issue is crucial for assessing the generalizability of findings between species. For example, the middle legs in stick insects do not show a prominent plane rotation as we observed here for *Drosophila.* However, protraction/retraction and levation/depression during stepping are performed by the ThCx and the CxTr, respectively (Büschges et al., 2008; Cruse et al., 2009) and the very same movements emerged from rotations about the ThCx-yaw and CxTr-yaw DOFs in our model. Hence, it is possible that morphological differences such as the orientation of joint rotational axes are the main cause for the dissimilarity in the movements of the middle legs in both species.

## 5 Conflict of Interest

The authors declare that the research was conducted in the absence of any commercial or financial relationships that could be construed as a potential conflict of interest.

## 6 Author Contributions

Conceptualization: M.H., A.B., T.B; Data curation: M.H.; Formal Analysis: M.H.; Funding acquisition: A.B.; Investigation: M.H., A.Bl.; Methodology: M.H., A.Bl., T.B.; Project administration: A.B.; Resources: A.B.; Software: M.H., T.B.; Supervision: A.B., T.B.; Validation: M.H.; Visualization: M.H.; Writing – original draft: M.H., T.B.; Writing – review & editing: M.H., A.Bl., T.B, A.B.

## 7 Funding

M.H. and A.B. are members of the “iBehave” network funded by the Ministry of Culture and Science of the State of North Rhine-Westphalia. T.B., A.Bl., and A.B. were supported by research grant C3NS in the international NeuroNex Program and herein funded by the Deutsche Forschungsgemeinschaft (DFG Bu857/15 and BL1355/6-1 [436258345]). The study additionally received funding from the PSI (ID: 20171469) for the beamline experiment.

## 8 Acknowledgements

We would like to Dr. Gesa Dinges for providing the specimen for the generation of the µCT data. Additional thanks go to Michael Dübbert and Mehrdad Ghanbari of the Electronics workshop of the Institute of Zoology, University of Cologne for their help with the motion capture setup. We also thank Fabian Jakobs for help in generating the training set for DeepLabCut and correcting tracking misdetections in the motion capture data. Parts of the text and the figures in this paper are reproduced from the doctoral dissertation of M.H. (Haustein, 2023). The here used motion capture data was partly published in two other studies (Goldsmith et al., 2022; Saltin et al., 2023).

## Notes

### Competing Interest Statement

The authors have declared no competing interest.

