## Supplemental Materials for "A leg model based on anatomical landmarks to study 3D joint kinematics of walking in *Drosophila melanogaster*"

[‡]Shared senior authorship

### 1 Supplementary Tables

**Supplemental-Table 1. Joint DOF angle constraints of the kinematic leg model**

| leg | joint | model DOFs | | | | | | orthogonalized model DOFs | | | | | |
|---|---|---|---|---|---|---|---|---|---|---|---|---|---|
| | | yaw (°) | | pitch (°) | | roll (°) | | yaw (°) | | pitch (°) | | roll (°) | |
| | | min | max | min | max | min | max | min | max | min | max | min | max |
| front | ThCx | -70 | 70 | -33 | 147 | -160 | 50 | -70 | 90 | -75 | 75 | -220 | 0 |
| | CxTr | -20 | 160 | -108 | 72 | -120 | 120 | -180 | 60 | -75 | 75 | -90 | 120 |
| | TrFe | -90 | 90 | -108 | 72 | -90 | 90 | -90 | 90 | -75 | 75 | -90 | 90 |
| | FeTi | -20 | 145 | -75 | 75 | -90 | 90 | -20 | 145 | -75 | 75 | -90 | 90 |
| | TiTar | -60 | 120 | -75 | 75 | n.a. | n.a. | -60 | 120 | -75 | 75 | n.a. | n.a. |
| middle | ThCx | -90 | 60 | -40 | 140 | -90 | 90 | -75 | 75 | -20 | 60 | -75 | 75 |
| | CxTr | -140 | 10 | -54 | 126 | -90 | 90 | -20 | 130 | -75 | 75 | -90 | 90 |
| | TrFe | -110 | 110 | -123 | 57 | -110 | 90 | -90 | 90 | -75 | 75 | -110 | 90 |
| | FeTi | 0 | 170 | -90 | 90 | -90 | 90 | -170 | 10 | -90 | 90 | -90 | 90 |
| | TiTar | -10 | 140 | -64 | 116 | n.a. | n.a. | 0 | 90 | -60 | 60 | n.a. | n.a. |
| hind | ThCx | -90 | 60 | -39 | 141 | -100 | 100 | -90 | 90 | -75 | 75 | -45 | 130 |
| | CxTr | -110 | 30 | -121 | 59 | -90 | 110 | -20 | 150 | -75 | 75 | -90 | 90 |
| | TrFe | -90 | 90 | -54 | 126 | -90 | 90 | -90 | 90 | -75 | 75 | -45 | 45 |
| | FeTi | 10 | 180 | -103 | 77 | -90 | 90 | -150 | 20 | -75 | 75 | -90 | 90 |
| | TiTar | -5 | 90 | -85 | 95 | n.a. | n.a. | -90 | 90 | -75 | 75 | n.a. | n.a. |

Note that values indicate constraints for joint DOFs of the right legs. The sign had to be inverted to obtain values for yaw and roll DOFs of the left legs, but not for pitch DOFs.

**Supplemental-Table 2. Area under curve (AUC) of the absolute model error time courses of different DOF configurations**

| model DOF configuration | | | | | | | front legs | | | | middle legs | | | | hind legs | | | |
|---|---|---|---|---|---|---|---|---|---|---|---|---|---|---|---|---|---|---|
| ThCx | | | CxTr | TrFe | | | swing | | stance | | swing | | stance | | swing | | stance | |
| yaw | pitch | roll | roll | yaw | pitch | roll | abs. | rel. | abs. | rel. | abs. | rel. | abs. | rel. | abs. | rel. | abs. | rel. |
| + | + | + | - | - | - | - | 47,662 | 1.00 | 39,152 | 1.00 | 12,845 | 1.00 | 10,899 | 1.00 | 13,761 | 1.00 | 13,645 | 1.00 |
| + | - | - | - | - | - | - | 166,357 | 3.49 | 110,492 | 2.82 | 16,602 | 1.29 | 15,134 | 1.39 | 25,547 | 1.86 | 30,590 | 2.24 |
| - | + | - | - | - | - | - | 105,713 | 2.22 | 87,040 | 2.22 | 29,892 | 2.33 | 24,263 | 2.23 | 36,196 | 2.63 | 33,323 | 2.44 |
| - | - | + | - | - | - | - | 170,984 | 3.59 | 115,093 | 2.94 | 27,021 | 2.10 | 26,029 | 2.39 | 41,122 | 2.99 | 49,478 | 3.63 |
| + | + | - | - | - | - | - | 82,128 | 1.72 | 64,836 | 1.66 | 15,672 | 1.22 | 13,551 | 1.24 | 17,295 | 1.26 | 17,630 | 1.29 |
| + | - | + | - | - | - | - | 160,174 | 3.36 | 103,566 | 2.65 | 14,967 | 1.17 | 13,708 | 1.26 | 23,989 | 1.74 | 28,815 | 2.11 |
| - | + | + | - | - | - | - | 95,289 | 2.00 | 79,632 | 2.03 | 15,834 | 1.23 | 13,560 | 1.24 | 23,728 | 1.72 | 24,947 | 1.83 |
| + | + | + | - | + | - | - | 27,291 | 0.57 | 25,770 | 0.66 | 12,792 | 0.93 | 10,793 | 0.99 | 11,585 | 0.84 | 10,688 | 0.78 |
| + | + | + | - | - | + | - | 43,826 | 0.92 | 34,417 | 0.88 | 10,994 | 0.86 | 9,173 | 0.84 | 13,501 | 0.98 | 13,297 | 0.97 |
| + | + | + | - | - | - | + | 21,692 | 0.46 | 21,592 | 0.55 | 12,283 | 0.96 | 10,563 | 0.97 | 12,300 | 0.89 | 10,571 | 0.77 |
| + | + | + | - | + | - | + | 19,833 | 0.42 | 20,004 | 0.51 | 12,800 | 1.00 | 10,837 | 0.99 | 12,790 | 0.93 | 11,280 | 0.83 |
| + | + | + | - | - | + | + | 21,474 | 0.45 | 20,537 | 0.52 | 10,537 | 0.82 | 8,828 | 0.81 | 12,258 | 0.89 | 10,434 | 0.76 |
| + | + | + | - | + | + | - | 26,987 | 0.57 | 24,624 | 0.63 | 11,009 | 0.86 | 9,314 | 0.85 | 11,856 | 0.86 | 10,582 | 0.78 |
| + | + | + | + | - | - | - | 22,412 | 0.47 | 22,772 | 0.58 | 12,009 | 0.93 | 10,240 | 0.94 | 12,779 | 0.93 | 11,216 | 0.82 |

Note that only DOFs that were different in the leg joint configurations are indicated. Abs. is the AUC of the mean model error time course for swing and stance phases. Rel. is the ratio between the AUC of the tested model and the reference model. Values for the reference model are given in the first row.
